# A highly contiguous genome assembly of *Brassica nigra* (BB) and revised nomenclature for the pseudochromosomes

**DOI:** 10.1101/2020.06.29.175869

**Authors:** Kumar Paritosh, Akshay Kumar Pradhan, Deepak Pental

## Abstract

*Brassica nigra* (BB), also called black mustard, is grown as a condiment crop in India. *B. nigra* represents the B genome of U’s triangle and is one of the progenitor species of *B. juncea* (AABB), an important oilseed crop of the Indian subcontinent. We report here a highly contiguous genome assembly of *B. nigra* variety Sangam. The genome assembly has been carried out using Oxford Nanopore long-read sequencing and optical mapping. The resulting chromosome-scale assembly is a significant improvement over the previous draft assemblies of *B. nigra*; five out of the eight pseudochromosomes were represented by one scaffold each. The assembled genome was annotated for the transposons, centromeric repeats, and genes. The *B. nigra* genome was compared with the recently available contiguous genome assemblies of *B. rapa* (AA), *B. oleracea* (CC), and *B. juncea* (AABB). Based on the maximum homology among the three diploid genomes of U’s triangle, we propose a new nomenclature for *B. nigra* pseudochromosomes, taking the *B. rapa* pseudochromosome nomenclature as the reference.

## 1. Introduction

Nagaharu^1^ based on his observations and preceding cytogenetic work^2^ proposed a model on the relationship of some of the cultivated Brassica species. The model, known as U’s triangle, described the relationship of three diploid species – *B. rapa* (Bra, AA, n=10), *B. nigra* (Bni, BB, n=8) and *B. oleracea* (Bol, CC, n=9) with three allopolyploid species – *B. juncea* (Bju, AABB, n=18), *B. napus* (Bna, AACC, n=19) and *B. carinata* (Bca, BBCC, n=17). Subsequent cytogenetic work on inter-specific and inter-generic hybrids between the Brassica species of the U’s triangle and other taxa in the tribe Brassiceae showed close relationships and the group was described as Brassica coenospecies^3,4^.

Since the early cytogenetic work, major insights have been gained into the evolution of the Brassica species based on the extent of nucleotide substitutions in the orthologous genes belonging to the nuclear^5^ and plastid genomes^6,7,8,9^, analysis of genome synteny using molecular markers^10,11^, *in situ* hybridizations^12^, and genome sequencing^13,14,15,16^. The most significant observation is that the three diploid species of the U’s triangle – *B. rapa, B. nigra, B. oleracea,* and the other diploid species belonging to the tribe Brassiceae have originated through genome triplication, referred to as the ***b*** event^5^. Genome triplication was followed by extensive chromosomal rearrangements leading to gene block reshuffling vis-à-vis the gene block order in *Arabidopsis thaliana* (At)^17,18^, and gene fractionation due to a differential loss of genes in the three constituent paleogenomes^19^. The diploid species of the tribe Brassiceae are, therefore, mesohexaploids. It is now accepted that tribe Brassiceae is defined by the ***b*** event; it is, however, not clear whether the ***b*** event happened once or more times. The presence of two plastid lineages^6,7,8,9^ points to a minimum of two independent ***b*** events^20^.

Genome assemblies of *B.rapa*^13^, *B. oleracea*^14^, *B. napus*^15^, and *B. juncea*^16^ were first reported using short-read Illumina sequencing. More recent assemblies of these species have used long-read sequencing technologies, either PacBio SMRT (single-molecule real-time) sequencing or Oxford Nanopore Technologies (ONT) ^21,22,23^. Scaffolding has been carried out with optical mapping and/or Hi-C technologies. The most extensive assembly of the B genome has been made available from our recent effort on the genome assembly of an oleiferous type of *B. juncea* variety Varuna with SMRT sequencing and optical mapping^23^.

We report here a highly contiguous genome assembly of *B. nigra* variety Sangam, a photoperiod insensitive, short-duration variety, grown under dryland conditions, and used as a seed condiment crop in India. The assembly has been carried out using Nanopore sequencing and optical mapping. Previously reported Illumina short-read sequences and a genetic map of *B. nigra*^23^ were used for error correction and assigning the contigs and scaffolds to the eight pseudochromosomes. We compared the structure of the B genome of *B. nigra* (BniB) with the genomes of *B. rapa*^21^ (BraA), *B. oleracea*^22^ (BolC), and also the B genome of *B. juncea*^23^. (BjuB). We propose a revised nomenclature for the *B. nigra* pseudochromosomes based on maximum homology between the A and B genome pseudochromosomes; the *B. rapa* A genome nomenclature being the reference as it was the first Brassica genome that was sequenced^13^.

## 2. Materials and methods

### 2.1. Plant material, genome size estimation, nanopore sequencing, optical mapping, and genome assembly

A DH (doubled haploid) line BnSDH-1 of *Brassica nigra* variety Sangam^23^ was used for genome sequencing and assembly. BnSDH-1 was maintained by bud pollination. For DNA isolation, BnSDH-1 seedlings were grown in a growth chamber maintained at 8 h light, 25° C / 16 h dark, 10° C cycle. DNA was isolated from the leaves of 10 d old seedlings; the harvested leaves were immediately frozen in liquid nitrogen. High molecular weight DNA was isolated from the leaf tissues by CTAB method^24^. For Nanopore sequencing, genomic DNA libraries were prepared using the ‘Ligation sequencing kit 1D’ following the manufacturer’s instructions (Oxford Nanopore). In brief, around 2 μg of high molecular weight DNA was repaired using the ‘NEBNext FFPE DNA Repair mix’ and the ‘Ultra ll End-prep Enzyme mix’; subsequently, the adapter mix was ligated to the repaired DNA using the ‘NEBNext Quick T4 DNA Ligase’. At the end of each step, DNA was cleaned with the ‘AMPure XP beads’ (Thermo Fisher Scientific). The quality and quantity of the DNA libraries were determined with a Nanodrop spectrophotometer. DNA libraries were sequenced on the MinION device using the MinION Flow Cells R 9.4.1 (Oxford Nanopore). Base-calling and quality filtering were carried out using Albacore software (https://github.com/Albacore). Previously generated^23^ Illumina short-read sequencing data (~100x coverage) of the line BnSDH-1 was used at various steps (described wherever used) of the new genome assembly. Approximately 40x Illumina PE (2×100 bp) data with a kmer length of 21 was used for the kmer frequency distribution analysis with Jellyfish v2.2.6^25^. The output histogram file was used to estimate the genome size of BnSDH-1 using the findGSE program^26^.

Raw Nanopore reads were assembled into contigs using the Canu assembler V1.6^27^ with the parameters ‘minRead length’ and ‘minOverlap length’ set at values of 1000 bp. The paired-end (PE) reads obtained earlier with Illumina sequencing (~100x coverage) were mapped on the assembled Nanopore contigs using BWA-MEM^28^, followed by error correction with the Pilon program^29^ in five iterative cycles. After each of the Pilon cycle, completeness of the corrected genome was ascertained with Benchmarking Universal Single Copy Orthologue (BUSCO) program (V4.0.5)^30^. OrthoDB v10 plant datasets were used as the reference for analyzing the completeness of the predicted genes.

Optical mapping was carried out following the protocols suggested by the manufacturer (Bionano Genomics). Leaf tissues from 7 d old seedlings were harvested and transferred to an ice-cold fixing solution. Nuclei were isolated using the ‘rotor-stator’ protocol (Bionano genomics, Document no: 30228) and the nuclear fraction was purified on a sucrose density gradient. The nuclei were embedded in 0.5 % w/v agarose followed by treatment with proteinase-K (Qiagen) for 2 h. Mapping was carried out with three different labeling reactions – two NLRS (Nicks, Labels, Repairs and Stains) and one DLS (Direct Label and Stain). For the two NLRS labeling reactions, agarose plugs were treated with BssSI and BspQI and the nicks were labeled with the ‘IrysPrep NLRS labeling kit’. In the DLS labeling reaction, DNA was recovered from the agarose plugs, suspended in TE buffer, and labeled with the ‘Bionano Prep DLS kit’. Mapping data were obtained from the labeled libraries on the Saphyr system (Bionano) using one lane for each library. Mapping and hybrid assemblies were performed using the Bionano Access software. A previously generated genetic map of *B. nigra*^23^, developed using an F_1_DH population from a cross of line BnSDH-1 × line 2782 was used for validating the scaffold level assemblies and assigning the scaffolds to the eight linkage groups (LGs) to constitute eight pseudochromosomes. Position of the GBS marker tags was determined on the scaffolds with a Blastn search analysis. A correlation plot of the physical and genetic position of the markers was developd to validate the integrity and quality of scaffolding. Scaffolds were positioned and oriented on each pseudochromosome based on the information obtained with the correlation plots.

#### 2.2. Transcriptome sequencing, gene, and transposon annotation

Illumina short-read based transcriptome sequencing of the line BnSDH-1 has been reported earlier^23^. The same protocol was used for transcriptome sequencing of the line 2782, an East European gene pool line of *B. nigra*. For transcriptome sequencing of the line BnSDH-1 on the PacBio platform, total RNA was isolated from the seedling, leaf, and developing inflorescence tissues using the ‘Spectrum plant total RNA kit’ (Sigma). The quality of the RNA was checked with Bioanalyzer 2100 using the ‘RNA 6000 Nano kit’ (Agilent). RNA samples with RIN values >7 were used for further analysis. Transcriptome sequencing was carried out on the pooled RNA. Three different libraries of the size range – 0.5 – 1 kb, 1 – 2 kb, and 2 – 6 kb were prepared using the ‘SMRTbell Template Prep kit’ and sequenced on a PacBio RS II sequencer. The raw sequences obtained from each of the three libraries were assembled separately using SMRT Analysis software (v1.4). Full-length non-chimeric sequences were used for clustering with ICE (Structure Clustering and Error Correction) algorithm; partial reads were used for polishing of the ICE generated consensus sequences. ORFs were predicted from the polished consensus sequences using the ANGEL software (https://github.com/PacificBiosciences/ANGEL).

Transposable elements (TEs) were identified in the genome assembly using the Repeatmodeler pipeline (http://www.repeatmasker.org/RepeatModeler/). A *de-novo* repeat library was developd using RECON, RepeatScout and Tandem Repeat Finder programs available in the Repeatmodeler pipeline, and NSEG (ftp://ftp.ncbi.nih.gov/pub/seg/nseg/) program. The developed TE library, along with the repbase database for At was used to predict TEs in the assembled genome using RepeatMasker (http://www.repeatmasker.org). Identified LTR sequences were validated by the LTR finder program^31^.

For gene annotation, repeat-masked genome assembly was used to predict the protein-coding genes with the Augustus program^32^ trained with 250 randomly selected *B. rapa* genes as the reference data set. The predicted genes were validated by a blast search against the Uniprot protein database (e value threshold < 1e-05). The predicted genes were validated by mapping the previously generated Illumina RNA-seq sequences^23^, and the RNA-seq and Iso-seq sequences generated in this study on the assembled genome. Illumina RNA-seq reads were mapped with STAR aligner^33^, and Iso-seq sequences were mapped using Minimap2 program^34^.

#### 2.3. Syntenic block identification and determination of gene fractionation patterns

Syntenic regions in the assembled genome were identified with the MCScanX program^35^. An all-against-all Blastp comparison was carried out between the *B. nigra* assembly and previously reported BraA^21^, BjuA and BjuB^23^, and At genome assemblies (e-value threshold 1e-05). The blastp output file was used along with the information of positions of each gene in all the genomes for synteny analysis. Parameters for the MCScanX were set as match_score: 50, match_size: 5, gap_penalty: −1, e-value: 1e-05, max_gaps: 25. Genes retained in each of the syntenic regions were calculated in a sliding window of 500 flanking genes at a given locus of At.

For divergence analysis, DNA sequences and the protein sequences of At genes and their orthologs in the BraA, BniB, and BjuB genomes were aligned with MUSCLE v3.8.31 software^36^. Poorly aligned regions were trimmed using GBLOCKS (v0.91)^37^ and PAL2NAL scripts^38^. A custom Perl script was used for the conversion of the aligned fasta format to the Phylip format. The Phylip files were converted into Newick format trees, and the Ks values were obtained using the PAML package^39^.

### 3. Results

#### 3.1. Genome sequencing and assembly

The *B. nigra* genome was reported to be ~591 Mb in size on the basis of nuclear DNA estimations with flow cytometry^16^. We estimated the size of *B. nigra* Sangam (line BnSDH-1) by using kmer frequency distribution of ~40x Illumina PE reads. The size of the genome was estimated to be ~522 Mb (Supplementary Fig. 1). Genome sequencing of the *B. nigra* line BnSDH-1 on the Nanopore MinION platform yielded a total of 8,778,822 reads with an N50 value of ~10 kb (Supplementary Table 1). The obtained long-reads provided ~100x coverage of the *B. nigra* genome if we consider the genome size to be ~522 Mb. The raw reads were assembled into 1,549 contigs with an N50 value of ~1.48 Mb using the Canu assembler (Table 1). The total size of the assembled contigs was ~515.4 Mb, covering ~ 98% of the *B. nigra* genome. Nanopore contigs were error-corrected with the ~100x Illumina PE^23^ reads using the Pilon program for five iterative cycles. A total of 124,464 nucleotide errors and 229,767 InDels were corrected. Most of the errors, predominantly present in the non-coding regions, could be identified and corrected in the first two cycles (Supplementary Fig. 2). The quality of the error-corrected contigs was ascertained after each cycle using BUSCO scores. At the end of the five correction cycles, 95.4% of the gene models were found to be complete.

**Table 1.**
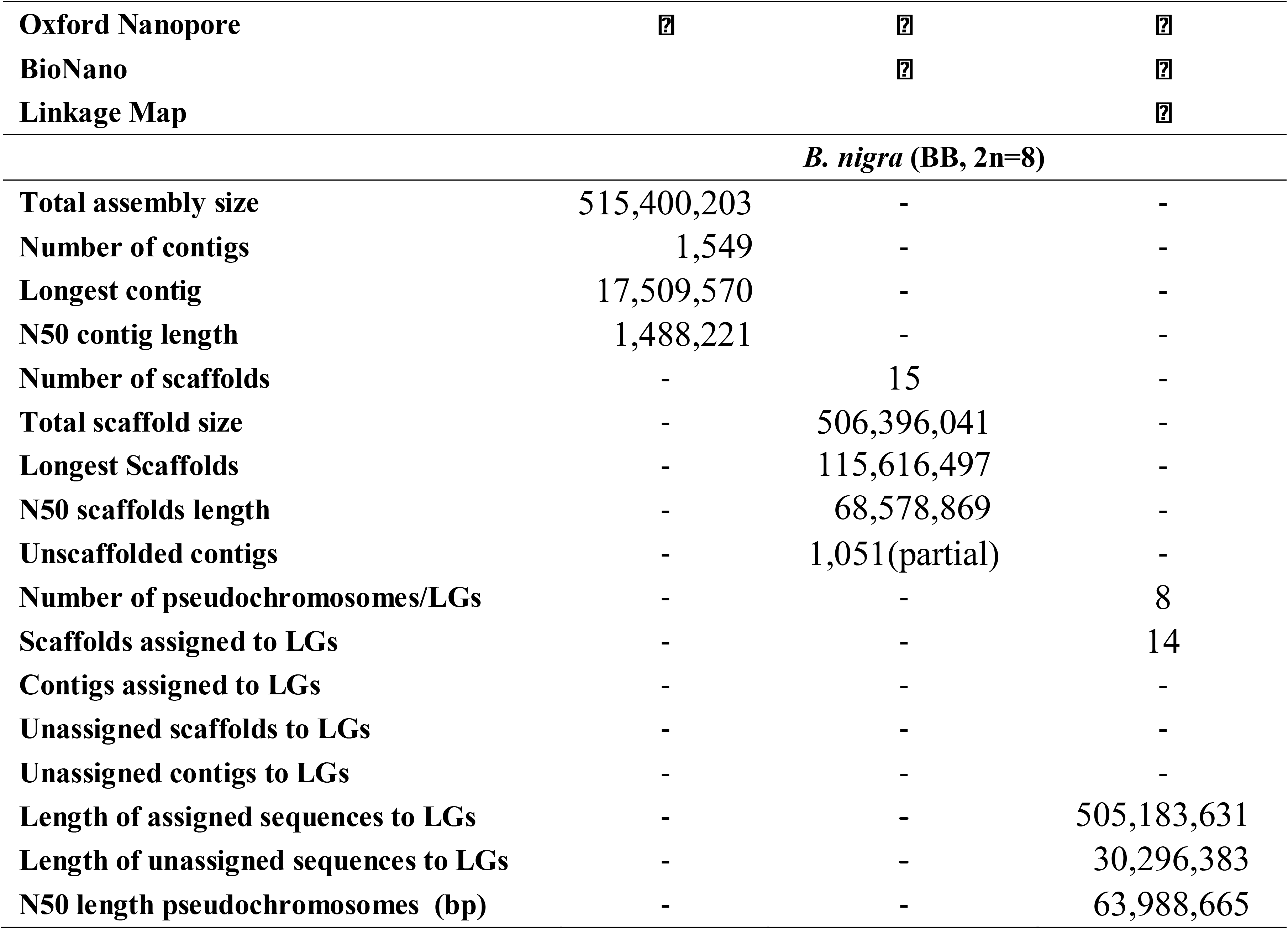
Genome assembly statistics of *B. nigra* variety Sangam.

Optical mapping was used for finding the misassemblies in the contigs and for assembling the contigs into scaffolds. Three different libraries, one with DLE labeling and two with NLRS labeling were used for optical mapping. Mapping data obtained from each one of the three reactions were assembled by hierarchical mapping protocol to develop a consensus map (CMAP). A total of 143 contigs were found to contain misassemblies, mostly due to the merger of some of the highly conserved syntenic regions. The misassembled contigs were curated by breaking such regions at the conflicting junctions and aligning these again with the CMAP in an iterative manner. The corrected contigs could be assembled into 15 scaffolds using the CMAP (Table 1). The total size of the scaffold level assembly was ~506.4 Mb, with an N50 value of ~68.6 Mb. A total of 1,051 unmapped sequence fragments, encompassing ~30.4 Mb of the genome with an N50 value of ~36.7 kb, remained unscaffolded.

A genetic map of *B. nigra*, with 2,723 markers^23^, was used to validate the integrity of the scaffolds and to assign these to the eight pseudochromosomes – BniB01 – BniB08 (Fig. 1). The genotyping by sequencing (GBS) based genetic markers were physically mapped on the scaffolds; no misassemblies were observed. Fourteen out of 15 scaffolds could be assembled into eight pseudochromosomes. Five out of the eight chromosomes were represented by a single scaffold each; the remaining three chromosomes consisted of two, three, and four scaffolds (Supplementary Table 2). One of the scaffolds was found to be unique as no genetic marker mapped on the scaffold; this scaffold consisted of the chloroplast genome of *B. nigra*. The size of the final *B. nigra* genome that could be assigned to the pseudochromosomes was ~505.18 Mb (~96.7% of the estimated genome size).

**Fig. 1.**
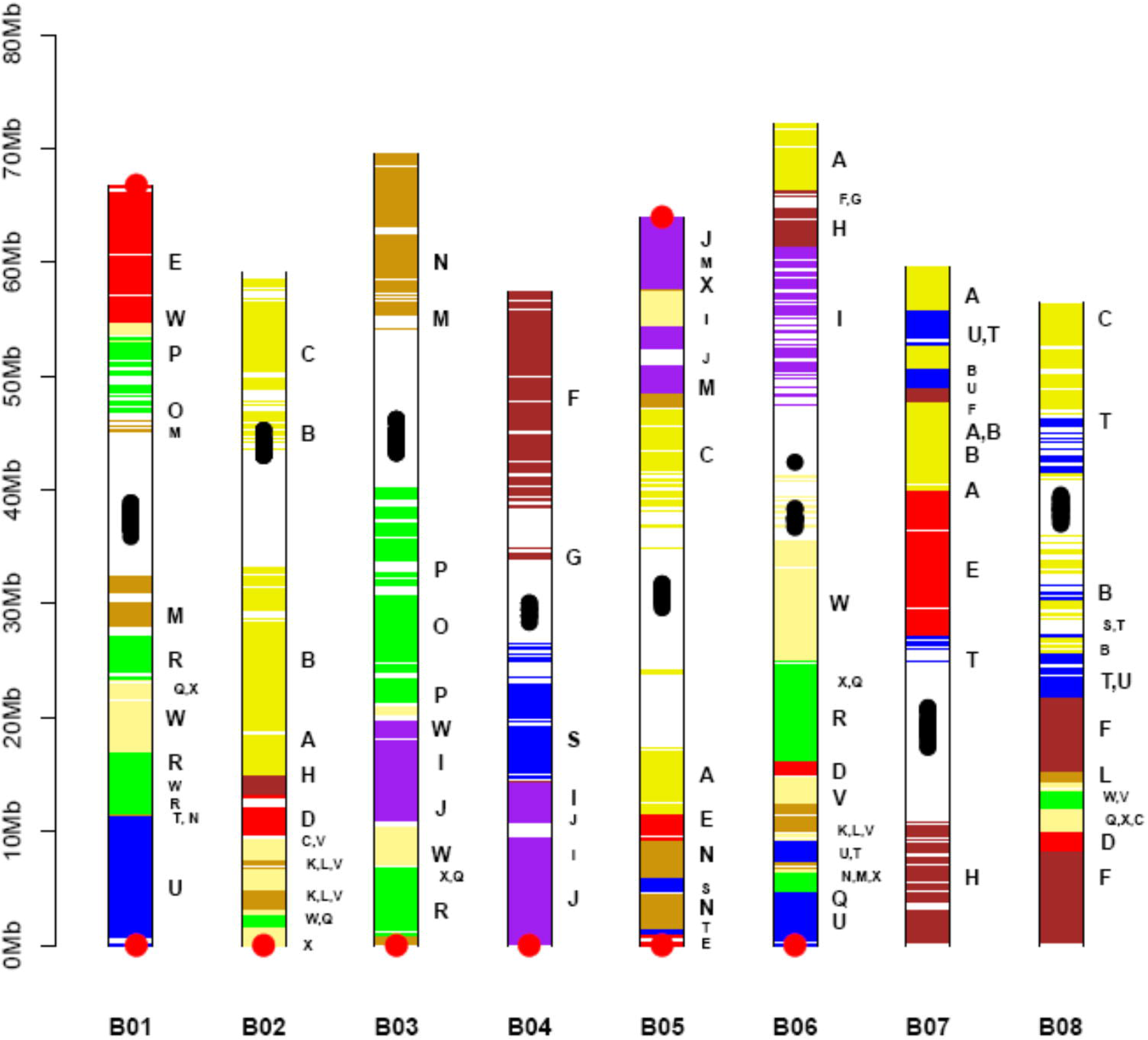
Graphic representation of the *Brassica nigra* pseudochromosomes. Each chromosome is represented by a vertical bar. Each horizontal bar represents a gene. Gene blocks have been identified on the basis of synteny with the *A. thaliana* gene blocks (A-X), as defined and color-coded by Schranz et al^17^. Centromeric repeats are represented as black dots and telomeric repeats as red dots. A new nomenclature has been given to the *B. nigra* pseudochromosomes on the basis of maximum gene-level collinearity with the *B. rapa* pseudochromosomes^21^.

#### 3.2. Genome annotation for repeat elements, centromeres, and genes

The assembled genome was annotated for the repeat elements, centromeric repeats, and genes. A *de-novo* prediction approach was used for the identification of the TEs. A repeat library was developed following the steps described in the Materials and methods section. *B. nigra* genome contained ~ 246 Mb (47.12 %) of repeat elements belonging to three broad categories – DNA transposons, retrotransposons, and other repeat elements. DNA transposons constituted ~31 Mb of the assembled genome; ~157 Mb of the genome was constituted of retrotransposons. LTR/Gypsy types were found to be the most predominant, ~103.1 Mb of the *B. nigra* genome; followed by ~43.6 Mb of LTR/Copia types (Supplementary Fig. 3). LTR/Copia types were found to be most abundant in the vicinity of the centromeric regions. Around 59 Mb of the repeat elements belonged to the unknown repeat category. We earlier carried out a study of the repeat elements constituting the centromeric regions in the B genome of *B. juncea*^23^. The centromere-specific repeats were identified as highly abundant kmers in the putative centromeric regions of the BjuB genome and were characterized for their sequences and their distribution (described in details in the reference 23); identical repeats were observed to constitute the *B. nigra* centromeric regions (Supplementary Fig. 4).

For gene annotation, the *B. nigra* pseudochromosome level assembly was repeat masked and used for gene prediction with the Augustus program^32^ trained with *B. rapa* gene content information. A total of 57,249 protein-coding genes were predicted in the *B. nigra* genome. The predicted genes were validated by comparing these with the non-redundant proteins in the UniProt reference database (TrEMBL); a total of 50,233 genes could be validated at an e-value threshold of 10^−5^. The predicted genes were further validated by Illumina RNA seq data obtained from the seedling, leaf, and young inflorescence tissues of the line BnSDH-1 and line 2782 (Supplementary File 1). A total of 39,946 genes could be validated by the transcriptome analysis. Transcriptome sequencing was also carried out on the PacBio platform (Supplementary File 1 for all the stats and description). A total of 15,368 full-length *B. nigra* genes were found in the Iso-seq analysis. The Iso-seq analysis validated 2,498 additional genes. Thus, a total of 42,444 genes, out of 57,249 predicted genes were validated by the transcriptome analysis of seedling, leaf, and developing inflorescence tissues (Supplementary Fig. 5).

#### 3.3. Gene block arrangement in *B. nigra*

The predicted 57,249 genes in *B. nigra* were checked for their syntenic gene block arrangements by comparisons with the gene block arrangements in the model crucifer At, and the two diploid species of the U’s triangle – *B. rapa* (AA)^21^, and *B. oleracea* (CC)^22^ with MCScanX. The *B. nigra* genome was divided into 24 gene blocks (A-X), identified in At^17^. Three syntenic regions were identified in the *B. nigra* genome for each gene block in At ( Supplementary Fig. 6).

Gene fractionation pattern was determined in each of the three *B. nigra* regions syntenic with each of the At gene blocks. Gene retention in the three syntenic regions in *B. nigra* was calculated by taking the number of genes present in the corresponding At gene block as a reference number. Based on the gene fractionation pattern, three sub-genomes were identified in the Bni genome – LF (Least Fragmented), MF1 (Moderately Fragmented), and MF2 (Most Fragmented) (Supplementary Fig. 6). In gene to gene comparison, the LF subgenome was found to contain 10,191 genes, MF1 8,822, and MF2 7,283 in comparison to a total of 19,091 genes present in the At genome. The three different syntenic regions with differential gene fractionation have been shown earlier to be a characteristic feature of the *B. rapa* and *B. oleracea* genomes^13,14^. The *B. nigra* genome and the B genome of *B. juncea* reported earlier^23^ show a similar pattern of gene fractionation in the three constituent paleogenomes.

The data on the physical position and the expression status of each predicted gene on the eight *B. nigra* pseudochromosomes Bni01 – Bni08 has been provided in the Supplementary Table 4. The data contains information on the ortholog of each At gene in the assembled *B. nigra* genome. We carried out the ortholog tagging of each gene of *B. nigra* and identified the nearest ortholog in *B. rapa* (BraA)^21^ and *B. juncea* (BjuB)^23^ genomes (Supplementary Table 4). A total of 24,799 genes were found to be BniB genome-specific; these could not be found in the syntenic regions of BraA and At genomes. Analysis of the transcriptome data showed 11,503 BniB genome-specific genes to be expressed.

#### 3.4. Comparison of B genome pseudochromosomes of *B. nigra* and *B. juncea*

We compared the B genome assembly of *B. nigra* line BnSDH-1 (BniB) with the B genome assembly of *B. juncea* line Varuna (BjuB)^23^ for the gene content, transposable elements, centromeric repeats, and syntenic regions based on gene collinearity. The repeat content in the BniB genome (~47.2%) was found to be similar to that in the BjuB genome (~51%). The LTR/Gypsy type transposons were the most abundant TEs followed by LTR/Copia types in both the genomes. The distribution of different types of TE elements was found to be similar in both the genomes.

Earlier six B genome-specific repeats were identified in the centromeric regions of the BjuB genome^23^. We found these repeats to be present in a similar manner in the centromeric regions of the *B. nigra* pseudochromosomes (Supplementary Fig. 4) and to be highly identical. In addition, CentBr1, CentBr2, and the other centromeric repeats reported to be present in the BraA, BolC, and BjuA genomes^13,14,23^ were absent in both the BjuB and BniB genomes. Our analysis indicates that the B genome has undergone a divergent evolutionary path than the A and C genomes in terms of the evolution of the centromeric repeats. The gene number estimation in the BniB genome (57,249) is very similar to the numbers predicted in the BjuB genome (57,084), suggesting no significant loss of genes in the B genome after allotetraploidization. Of a total of 22,498 B genome-specific genes identified in the BjuB genome, 19,175 genes were also detected in the BniB genome.

We compared the overall genome architecture of the BniB and BjuB genomes by MCScanX based analysis. Orthologous genes were identified as the syntenic gene pairs having the least Ks value amongst all the possible combinations. The homologous gene pairs between the two B genomes were plotted using the Synmap analysis^40^. Very high collinearity was observed between the BniB and the BjuB pseudochromosomes (Fig. 2). An inversion was observed in each of the three pseudochromosomes – BniB01, BniB04, and BniB08 vis-à-vis the corresponding BjuB pseudochromosomes. The inversions in the BniB01 and BniB08 pseudochromsomes were found to be intra-block inversions in the U and F gene blocks, respectively. An inter-paleogenome non-contiguous gene block association^23^ J_MF1_-I_MF1_-S_MF2_-S_LF_ observed in BjuB04 and shared with BraA04 and BolC04 was found to be J_MF1_-I_MF1_-J_MF1_-I_MF1_-S_MF2_-S_LF_ in BniB04. This new gene block association in BniB04 is due to an inversion in the J_MF1_-I_MF1._ This inversion seems to be specific to the sequenced Sangam genome. It can be concluded that the progenitor B genome of *B. juncea* did not contain all the three inversions.

**Fig. 2.**
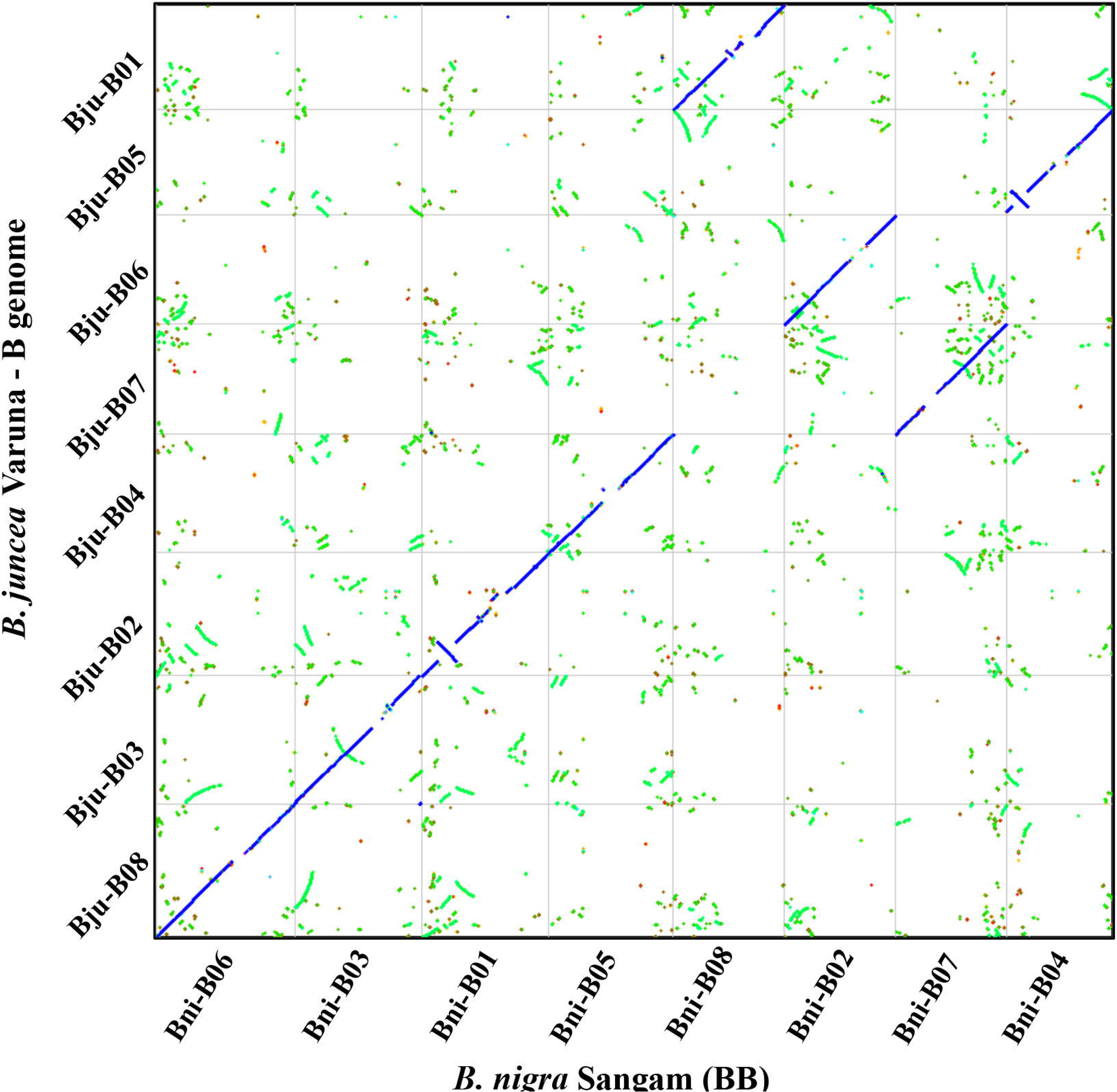
Comparison of *B. nigra* (BniB) pseudochromosomes with *B. juncea* B genome (BjuB) pseudochromosomes. The comparison was carried out with the Synfind program available at the CoGe website. Gene pairs with the least Ks value were identified as orthologous genes between the two genomes. Strictly orthologous genes have been denoted as blue dots, other syntenic regions are shown with the green dots. Very high gene collinearity was observed between the two B genomes, except for the three inversions in the *B. nigra* pseudochromosomes - BniB01, BniB04, and BniB08. Centromeric regions are devoid of genes and therefore, recognized as gaps. The nomenclature of the Bni pseudochromosomes is according to the new nomenclature, the BjuB pseudochromosome nomenclature is following Panjabi et al^11^.

#### 3.5. New nomenclature for *B. nigra* pseudochromosomes

Highly contiguous pseudochromosome level assemblies have been available for *B. rapa* (BraA)^21^, and *B. oleracea* (BolC)^22^; such an assembly is now available for *B. nigra* (BniB) allowing a chromosome level homology analysis. We carried out such an analysis for the BraA and BniB pseudochromosomes keeping the nomenclature given to the BraA^13^ pseudochromosomes as settled as it was the first sequenced genome from the U’s triangle. Each assembled pseudochromosome of *B. nigra* showed homology with more than one pseudochromosome of *B. rapa* (Fig. 3, Supplementary Fig. 7). The size of the genomic stretches from the Bra pseudochromosomes showing homology with different Bni pseudochromosomes was calculated (Table 2). Each BniB pseudochromosome was given the number of the BraA pseudochromosome with which it shared maximum homology (except pseudochromosome BniB02). As *B. nigra* has eight chromosomes against ten in *B. rapa*, homology with BraA09 and BraA10 was not taken into consideration. The new nomenclature is Version 3.

**Fig. 3.**
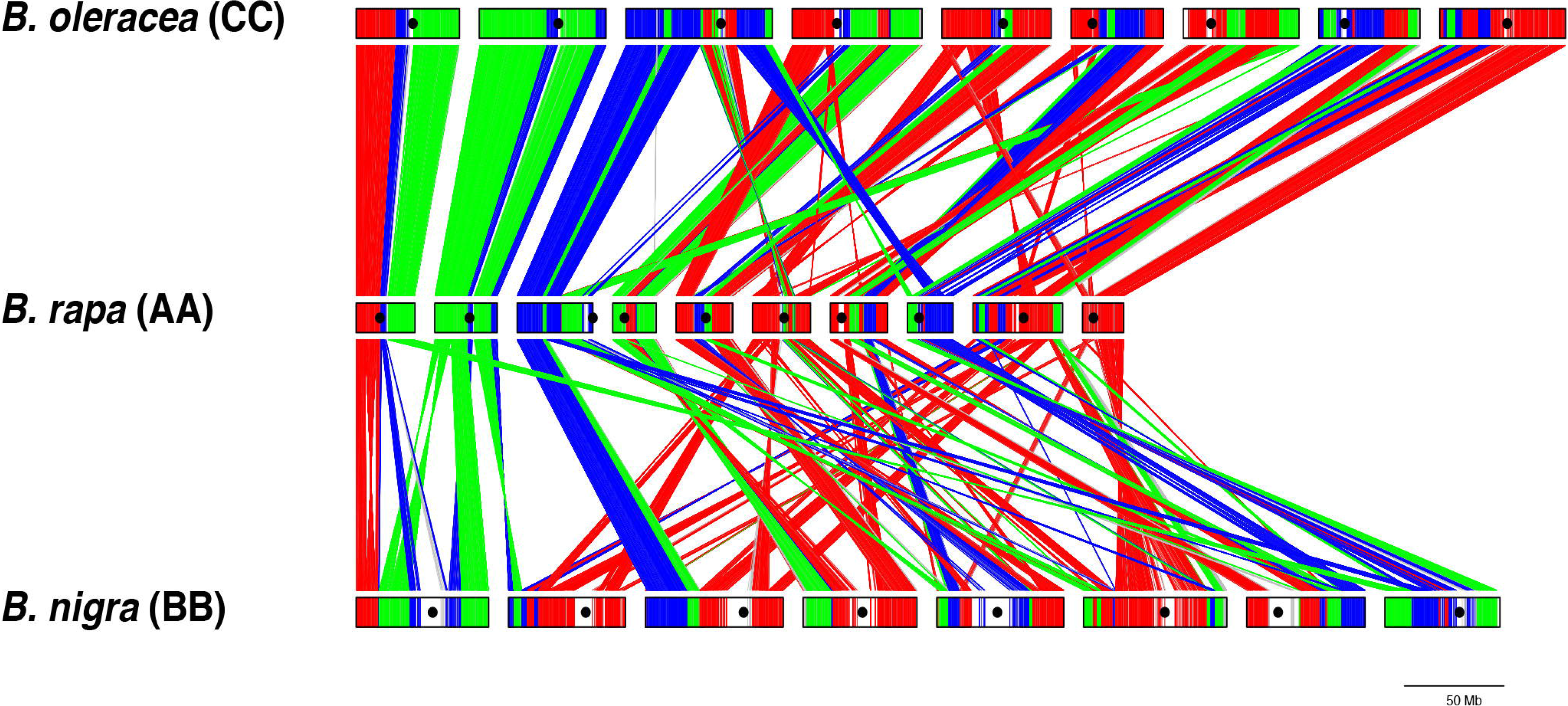
Comparative gene block arrangements in *B. rapa*^21^, *B. nigra* (this study), and *B. oleracea*^22^. All the three assemblies are with long-read sequences. The LF, MF1 and MF2 paleogenomes present in the A, B and C genomes have been represented by red, green and blue colors, respectively. The A and C genomes show more similarity in gene block arrangements, whereas the B genome has divergent arrangements.

**Table 2.**
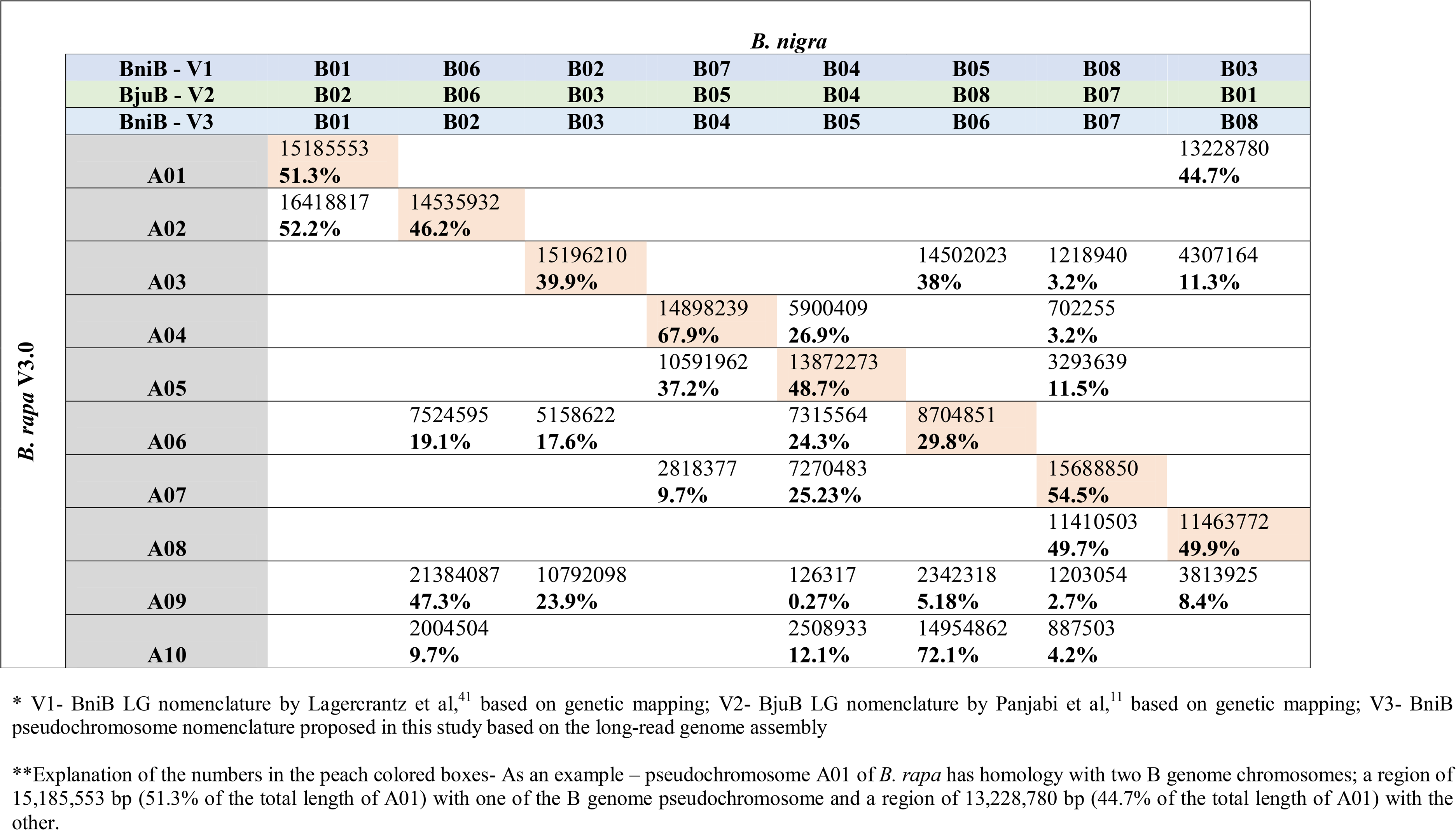
The size of the genomic stretches from the *B. rapa* pseudochromosomes showing gene collinearity-based homology with different *B. nigra* pseudochromosomes.

### 4. Discussion

The current nomenclature (Version 1) for the *B. nigra* LGs, recommended by the internationally agreed standard (http://www.brassica.info), is based on some early work on the comparative genetic mapping between At and *B. nigra*^41^. A total of 160 DNA fragments from the At genome, mostly anonymous and some cDNA fragments of known genes, were used as RFLP markers. We carried out a more extensive mapping work on the A and B genomes of *B. juncea* using intron length polymorphism (IP) markers derived from the At genome^11^. This allowed a more extensive comparative genetic mapping between the A and the B genomes of *B. juncea* vis-a-vis the gene block organization in the At genome. A different nomenclature (Version 2) was suggested for the BjuB genome LGs based on the extent of homology with the BjuA LGs. This nomenclature was supported by genetic mapping in *B. juncea* using RNAseq based SNP markers^42^.

While Version 1 and Version 2 are based on genetic mapping, Version 3 is based on gene collinearity and is, therefore, more accurate (Table 2). Version 1, due to low marker density is the most inaccurate. Version 1-B02 and B05 have no homologous regions with BraA02 and BraA05 chromosomes, respectively. Version 2 is more accurate; however, Version 2 - B08 has no homology with BraA08. The inter-paleogenome non-contiguous gene block association J_MF1_-I_MF1_-S_MF2_-S_LF,_ which is evidence for a common origin of the A, B, and C genomes^23^, is only accounted for in Version 3. We propose that Version 3 nomenclature be accepted by the Brassica community both for *B. nigra* and *B. juncea* pseudochromosomes, as it reflects gene-to-gene based homology between the genomes of the species of the U’s triangle.

*B. nigra* germplasm could be an important source for some of the major diseases afflicting the more extensively cultivated Brassica species. So far extensive efforts have been devoted to the transfer of resistance to the blackleg disease (causal organism *Leptosphaeria maculans*) from *B. nigra* to *B. napus*^43^. While the chromosomes of *B. nigra* containing resistance were identified in the chromosome addition lines^44^, actual introgression has been difficult due to limited pairing between the B, and the A and C genome chromosomes^45^. Genome assemblies of the B genome in *B. nigra* and *B. juncea* have shown a very divergent chromosomal organization between the B, and A/C genomes. Genetic exchanges may also be limited due to a strong mechanism in the B genome for suppression of pairing between the homeologous chromosomes^46,47^.

*B. nigra* genome assembly reported here is an improvement over the previous *B. nigra* assemblies based on short-read sequencing^16,48^. The long-read Nanopore sequencing and optical mapping have provided highly contiguous genomes assembly, with five of the eight pseudochromosomes represented by a single scaffold. Centromeric and telomeric regions could also be identified.

We have compared the *B. nigra* (BniB) genome assembly reported in this study with the B genome of *B. juncea* (BjuB) assembled with SMRT sequencing and optical mapping^23^, and shown that the two genomes are collinear in gene arrangement, and have similar gene content and centromeric structures. We have earlier shown that the A genome of *B. juncea* (BjuA)^23^ is similar to the *B. rapa* (BraA)^21^ genome. The success of the natural allotetraploid *B. juncea* was therefore based on immediate stability due to suppression of homoeologous pairing between the A and the B genome as has been suggested in some of the early cytogenetic studies^46,47^. However, high collinearity between BniB and BjuB genomes would allow the use of *B. nigra* germplasm for broadening the genetic base of *B. juncea* and transfer of disease resistance and other traits from *B. nigra* to *B. juncea*. As an example, *B. nigra* line 2782 is resistant to a number of isolates of oomycete pathogen *Albugo candida* and can be a useful source of resistance for the susceptible Indian gene pool lines of *B. juncea*^49^.

We have suggested a new nomenclature for the *B. nigra* LGs/chromosomes. The nomenclature currently in use does not follow any structural or evolutionary relationship with the other Brassica species of the U’s triangle or At. Any nomenclature should reflect some evolutionary relationships. The new nomenclature reflects the extent of homology between the B genome and the A and C genomes. We propose that the suggested nomenclature for the B genome LGs/chromosomes, taking the BraA chromosome nomenclature as settled, be accepted by the Brassica researcher community.

## Supporting information

Supplementary Tables B. nigra

Supplementary Figures B. nigra

Supplementary File 1 B. nigra

Supplementary Table 4 B. nigra

## Acknowledgments

The work was supported by the Department of Biotechnology (DBT), Government of India through two different grants – Centre of Excellence (Grant no.- BT/01/COE/08/06-II), and DBT-UDSC Partnership Centre on Genetic Manipulation of Brassicas (Grant no.- BT/01/NDDB/UDSC/2016). DP acknowledges support by a J C Bose Fellowship from the Department of Science and Technology (DST) and by the Council of Scientific and Industrial Research (CSIR) as a Distinguished Scientist.

## Authors contributions

KP carried out the genome assembly, gene annotation, and the other bioinformatics analysis; KP and DP wrote the manuscript, AKP and DP supervised the study. All the authors read and approved the final manuscript.

## Data availability

*B. nigra* genome and transcriptome sequences have been deposited under bioproject PRJNA324621 and PRJNA642332.

## Code availability

Codes used in the manuscript are available on request.

