## Supplementary Tables B. nigra for "A highly contiguous genome assembly of *Brassica nigra* (BB) and revised nomenclature for the pseudochromosomes"

**Supplementary Table 1**. ***Brassica nigra* variety Sangam raw sequencing data obtained with Oxford Nanopore machine**

| **PacBio raw reads** | **stats** |
| --- | --- |
| Number of reads | 8,778,822 |
| Total size of reads (bp) | 52,227,562,135 |
| Longest read | 182,926 |
| Number of reads > 1K nt | 7,608,368 |
| Number of reads > 10K nt | 1,558,964 |
| Number of reads > 100K nt | 136 |
| Mean read size | 5,949 |
| Median read size | 3,816 |
| N50 read length | 10,090 |
| L50 read count | 1,536,070 |

**Supplementary Table 2**. **Position and number of the assembled scaffolds on each of the eight pseudochromosomes of *Brassica nigra***

| **S. No** | **Chromosome** | **Number of scaffolds/**  **contigs** | **Scaffolds name** | **Length (bp)** | **Features** |
| --- | --- | --- | --- | --- | --- |
| 1 | BniB01 | 1 | Scaffold_4a | 56,519,733 | Centromere |
| 2 | BniB02 | 3 | Scaffold_7 | 48,531,046 | Telomere, Centromere |
|  |  |  | Scaffold_100000100016 | 203,637 |  |
|  |  |  | Scaffold_8 | 18,041,172 | Telomere |
| 3 | BniB03 | 2 | Scaffold_5 | 68,578,869 | Telomere, Centromere |
|  |  |  | Scaffold_12 | 899,017 |  |
| 4 | BniB04 | 4 | Scaffold_13 | 784,688 | Telomere |
|  |  |  | Scaffold_9 | 15,344,140 |  |
|  |  |  | Scaffold_10 | 4,249,561 |  |
|  |  |  | Scaffold_3 | 43,609,976 | Telomere, Centromere |
| 5 | BniB05 | 1 | Scaffold_6 | 57,438,084 | Telomere, Centromere |
| 6 | BniB06 | 1 | Scaffold_4b | 59,096,763 | Telomere, Centromere |
| 7 | BniB07 | 1 | Scaffold_1 | 59,653,443 | Centromere |
| 8 | BniB08 | 1 | Scaffold_2 | 72,232,902 | Telomere, Centromere |
|  | Other scaffolds | 1 | Scaffold_200000101 | 498,801 | Chloroplast |

**Supplementary Table 3. *Brassica nigra* Sangam genome – types of TEs and other repeats**

|  | **TE type** | **Repeat elements** | **Intact copies** | **Length** | **% genome coverage** |
| --- | --- | --- | --- | --- | --- |
| **Class I: DNA transposon** | DNA/CMC-EnSpm | 12933 | 11866 | 8896721 | **1.77** |
|  | DNA/Crypton-S | 1916 | 1701 | 690761 | **0.14** |
|  | DNA/hAT | 2601 | 2358 | 1270894 | **0.25** |
|  | DNA/hAT-Ac | 16363 | 15067 | 5949507 | **1.18** |
|  | DNA/hAT-Charlie | 863 | 849 | 482279 | **0.1** |
|  | DNA/hAT-Tag1 | 5442 | 5019 | 1509057 | **0.30** |
|  | DNA/hAT-Tip100 | 994 | 861 | 407834 | **0.08** |
|  | DNA/IS3EU | 296 | 286 | 107212 | **0.02** |
|  | DNA/Maverick | 157 | 155 | 52930 | **0.01** |
|  | DNA/Merlin | 44 | 33 | 11427 | **0.002** |
|  | DNA/MuLE-MuDR | 6156 | 5030 | 4456749 | **0.88** |
|  | DNA/PIF-Harbinger | 7263 | 6632 | 2467504 | **0.49** |
|  | DNA/RC | 4419 | 3526 | 1981015 | **0.39** |
|  | DNA/TcMar-Pogo | 3289 | 3063 | 756166 | **0.15** |
|  | DNA/TcMar-Stowaway | 7037 | 6912 | 1451632 | **0.29** |
|  | DNA/Zisupton | 1467 | 1304 | 412805 | **0.082** |
| **Subtotal** |  | **71240** | **64662** | **30904493** | **6.13** |
| **Class II: Retrotransposon** | LINE/L1 | 15495 | 14134 | 8975037 | **1.78** |
|  | LINE/Penelope | 923 | 879 | 295226 | **0.059** |
|  | LTR/Cassandra | 1560 | 1516 | 584873 | **0.12** |
|  | LTR/Caulimovirus | 987 | 947 | 605419 | **0.12** |
|  | LTR/Copia | 35484 | 29662 | 43630011 | **8.66** |
|  | LTR/DIRS | 233 | 230 | 76735 | **0.01** |
|  | LTR/ERV1 | 52 | 33 | 22473 | **0.005** |
|  | LTR/Gypsy | 65586 | 57202 | 103103094 | **20.46** |
|  | LTR/Pao | 162 | 134 | 31273 | **0.006** |
|  | SINE? | 124 | 124 | 7320 | **0.001** |
|  | SINE/tRNA | 3718 | 3687 | 562360 | **0.11** |
| **Subtotal** |  | **124324** | **108548** | **157893821** | **31.33** |
| **Other Repeats** | **Satellites** | **557** | **NA** | **164157** | **0.26** |
|  | **Simple repeats** | **1591** | **NA** | **577409** | **0.92** |
|  | **Unknown** | **160773** | **NA** | **59057277** | **9.38** |

**Supplementary Table 4. Genes predicted on different pseudochromosomes of *Brassica nigra* Sangam and their orthologs in *Arabidopsis thaliana* (along with their respective gene blocks) and *B. juncea* Varuna B genome (BjuB) and *B. rapa* Chiifu V3.0 (BraA) genome.** Column A – gene blocks as identified in *A. thaliana*; Column B – *A. thaliana* orthologs with gene id; Column C – paleogenome of *B. nigra* to which the gene belongs; Column D – predicted *B. nigra* gene id; Column E – physical position of the genes on the pseudochromosomes; Column F – expression status of the predicted *B. nigra* genes (“Expressed” means that the gene was found in the transcriptome analysis in this study or other studies described in Supplementary File 1, “Not expressed” represents – an expression not found); Column G – *B. juncea* B genomes (BjuB) orthologs gene id; Column H –*B. rapa* V3.0 (BraA) orthologs with the gene id.
