## Supplementary Figures B. nigra for "A highly contiguous genome assembly of *Brassica nigra* (BB) and revised nomenclature for the pseudochromosomes"

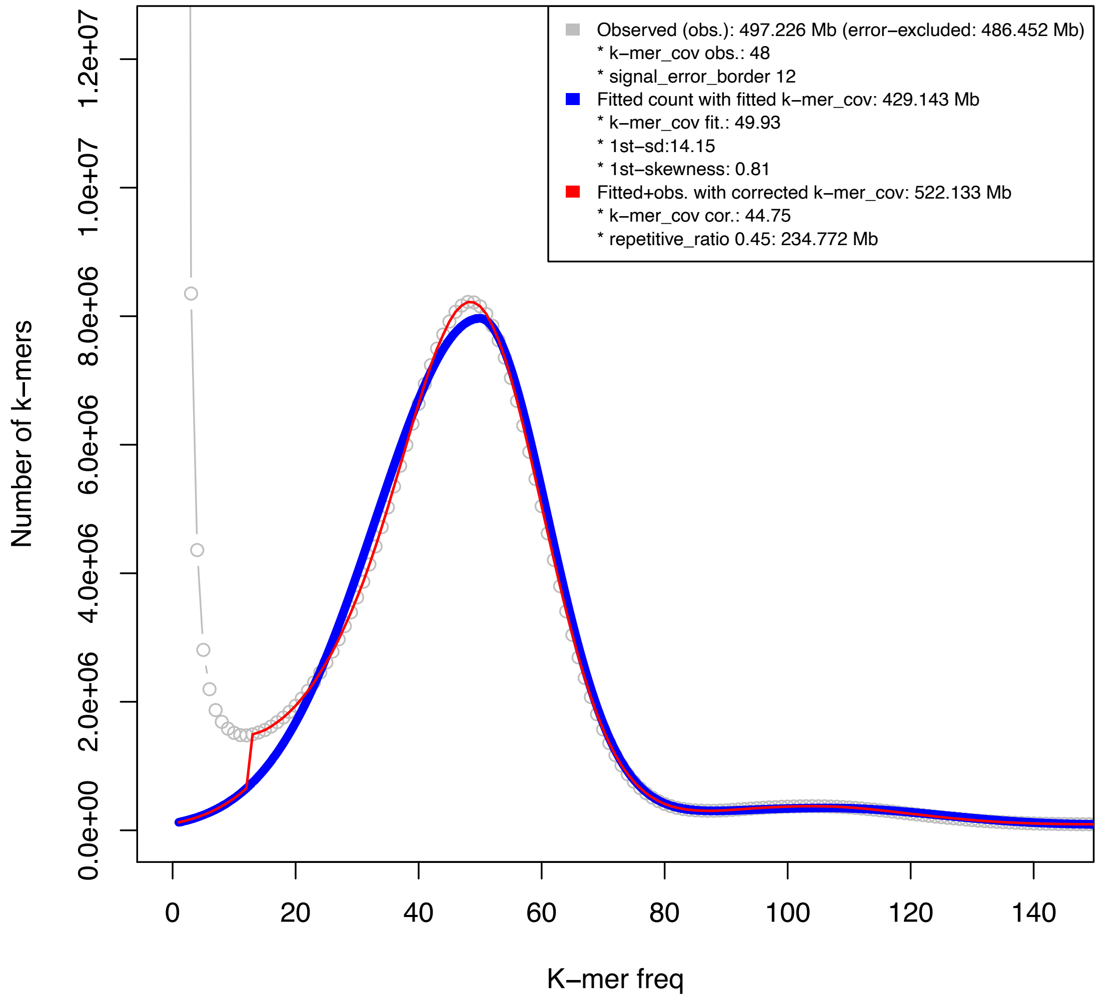


**Supplementary Fig. 1. K-mer frequency distribution in ~40x PE Illumina reads of *B. nigra* line BnSDH-1.** The frequency of the kmers of length 21bp was calculated and used to estimate the genome size with FindGSE program. The *B. nigra* genome was estimated to be ~522.13 Mb in size.

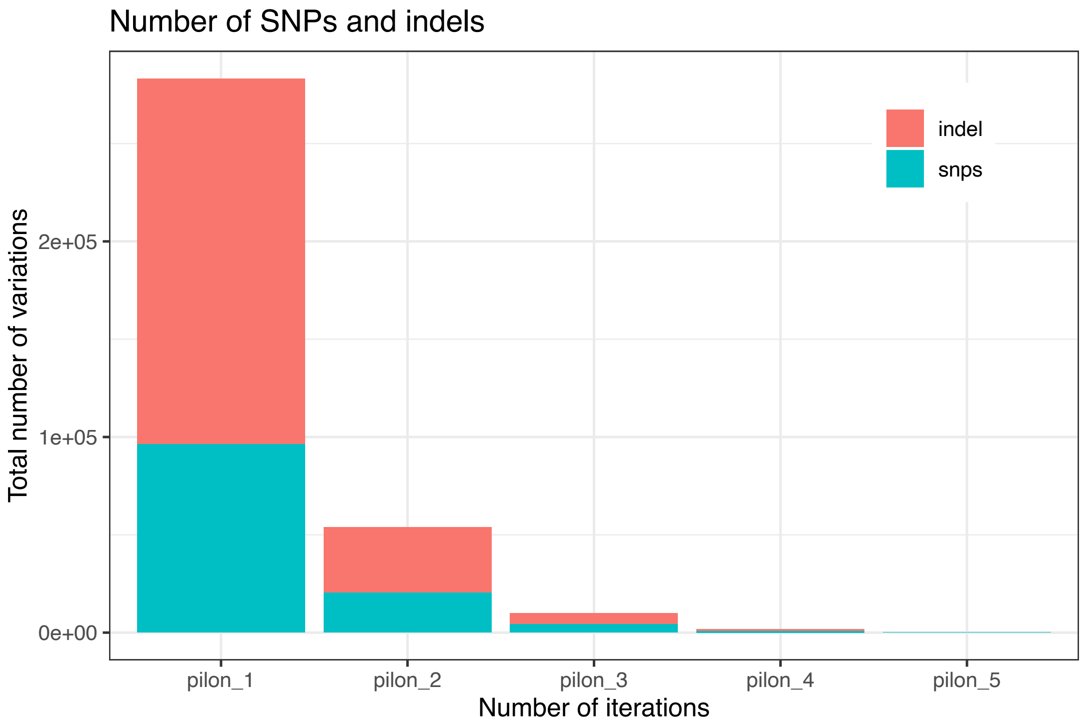


**Supplementary Fig. 2. Correction of *B. nigra* Nanopore assembly contigs with ~100x Illumina short-reads using the Pilon program.** Five rounds of Pilon based corrections of SNPs and InDels were carried out iteratively. Most of the errors were identified and corrected in the first two cycles.

### Distribution of Transposable elements in *B. nigra* genome

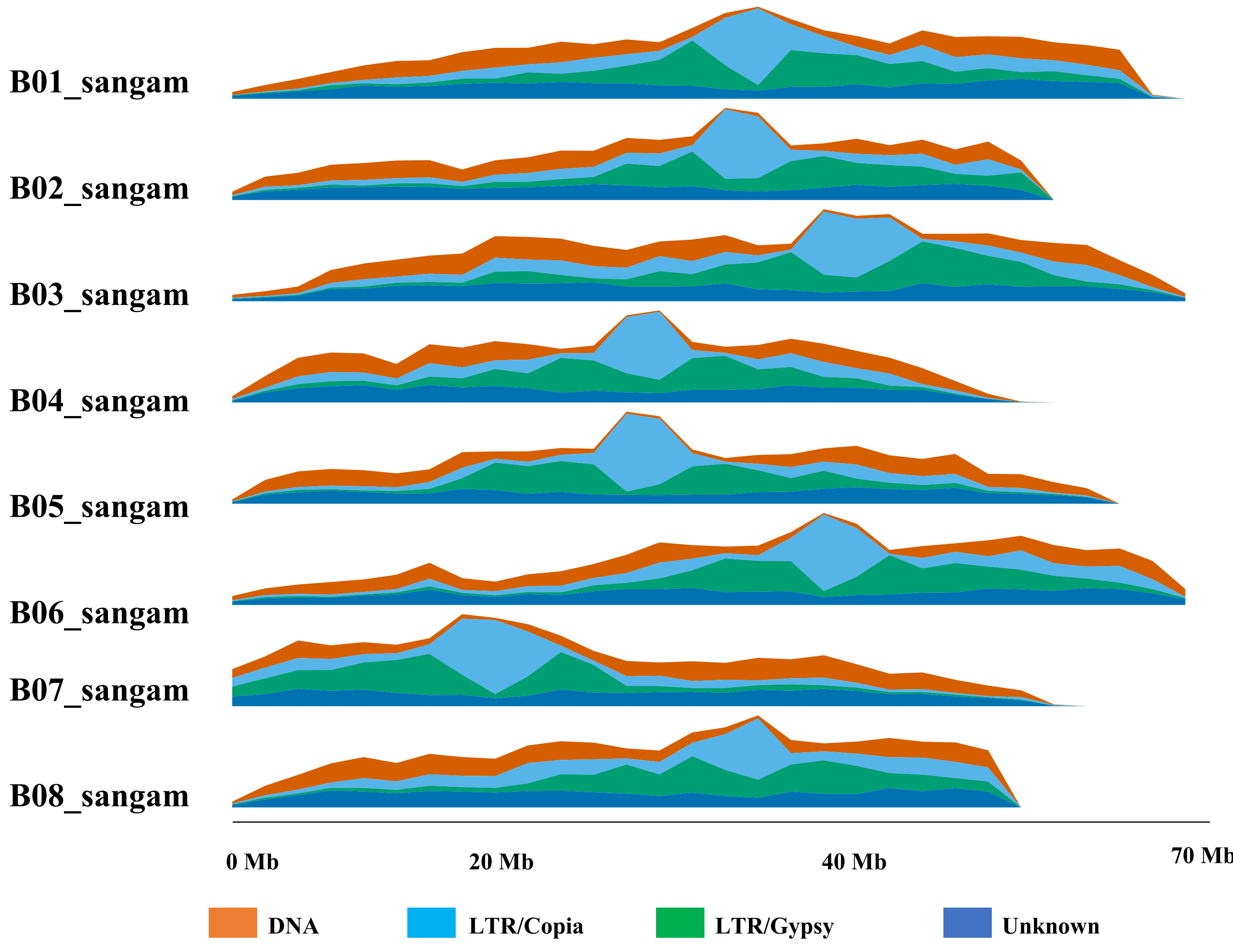


**Supplementary Fig. 3. Distribution of different TEs on the *B. nigra* pseudochromosomes.** LTR/Copia and LTR/Gypsy type transposable elements are the most abundant TEs. Centromeric regions show a much higher content of LTR/Copia TEs.

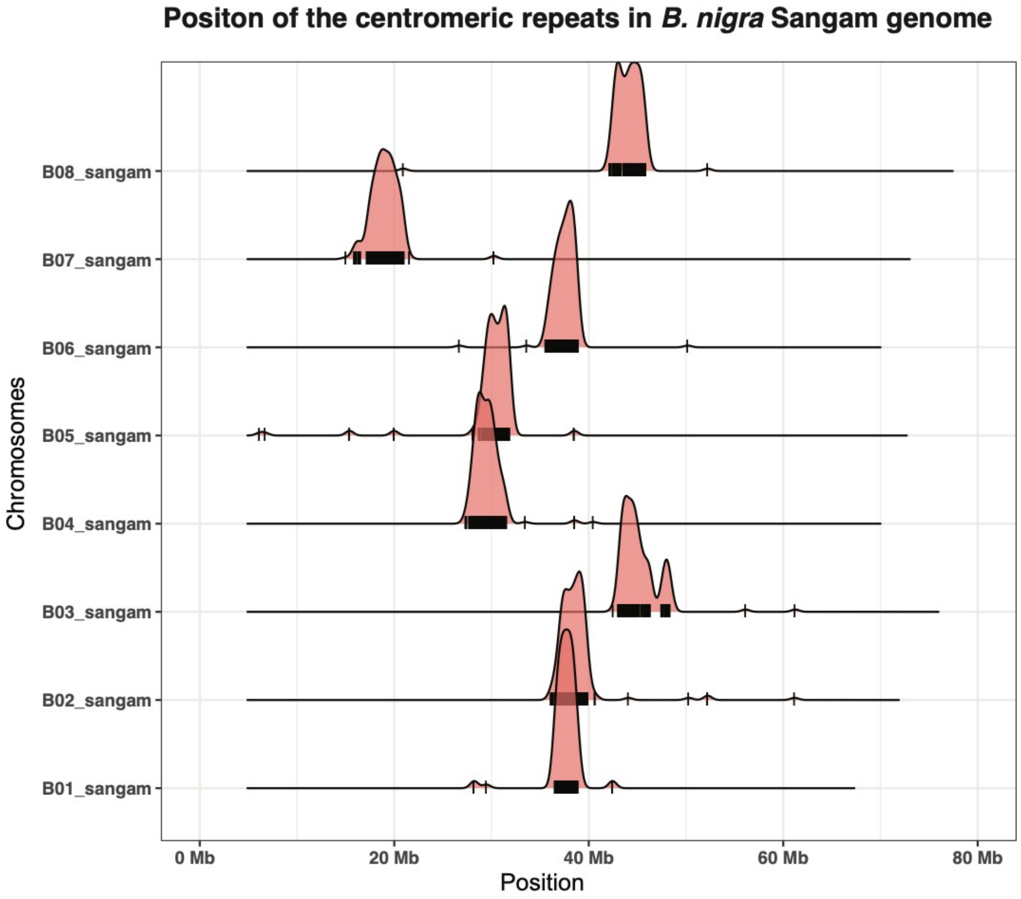


**Supplementary Fig. 4. Distribution of the B genome-specific centromeric repeats on the eight pseudochromosomes of *B. nigra*.** The earlier described six unique repeat sequences in the B genome of *B. juncea*[23] were found to constitute the centromeric regions of the *B. nigra* pseudochromosomes. The position of the centromeric repeats on the pseudochromosomes has been shown by horizontal bars; the vertical curve represents the cumulative number of the predicted centromeric repeats.

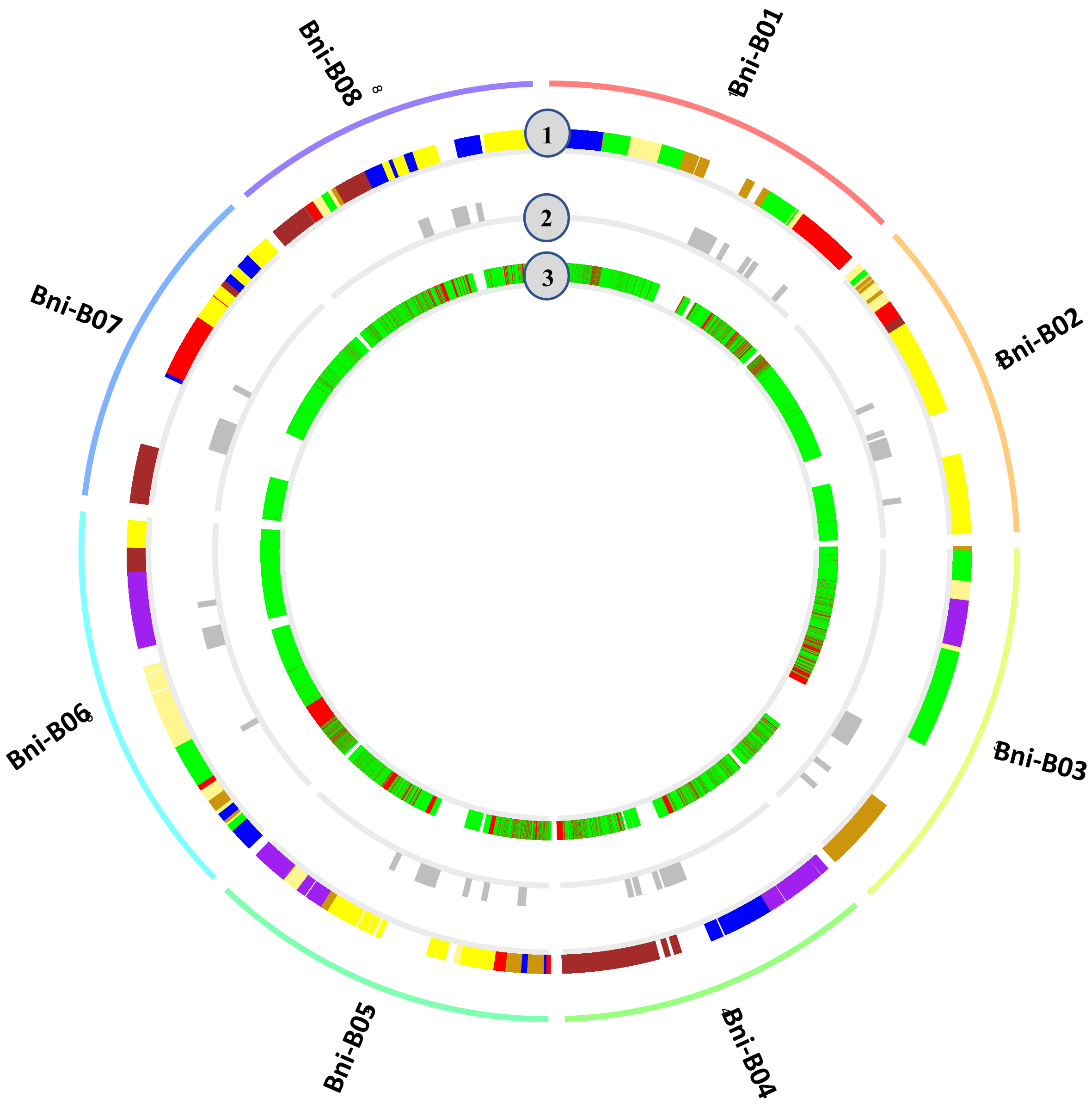


**Supplementary Fig. 5. Genes identified in the transcriptome of leaf, stem, and developing inflorescence of *B. nigra*.** Genes present on the eight pseudochromosomes of *B. nigra* are represented in track 1; each gene is represented by a line and colored based on the gene bock it belongs to, following Schranz et al[17]. Track 2 represents the position of the centromeric repeats in each of the *B. nigra* pseudochromosomes. Track 3 represents the position of the genes found to be expressed in the transcriptome sequencing data; each expressing gene has been represented by a green line; genes non-expressing in the transcriptome study are represented by red lines.

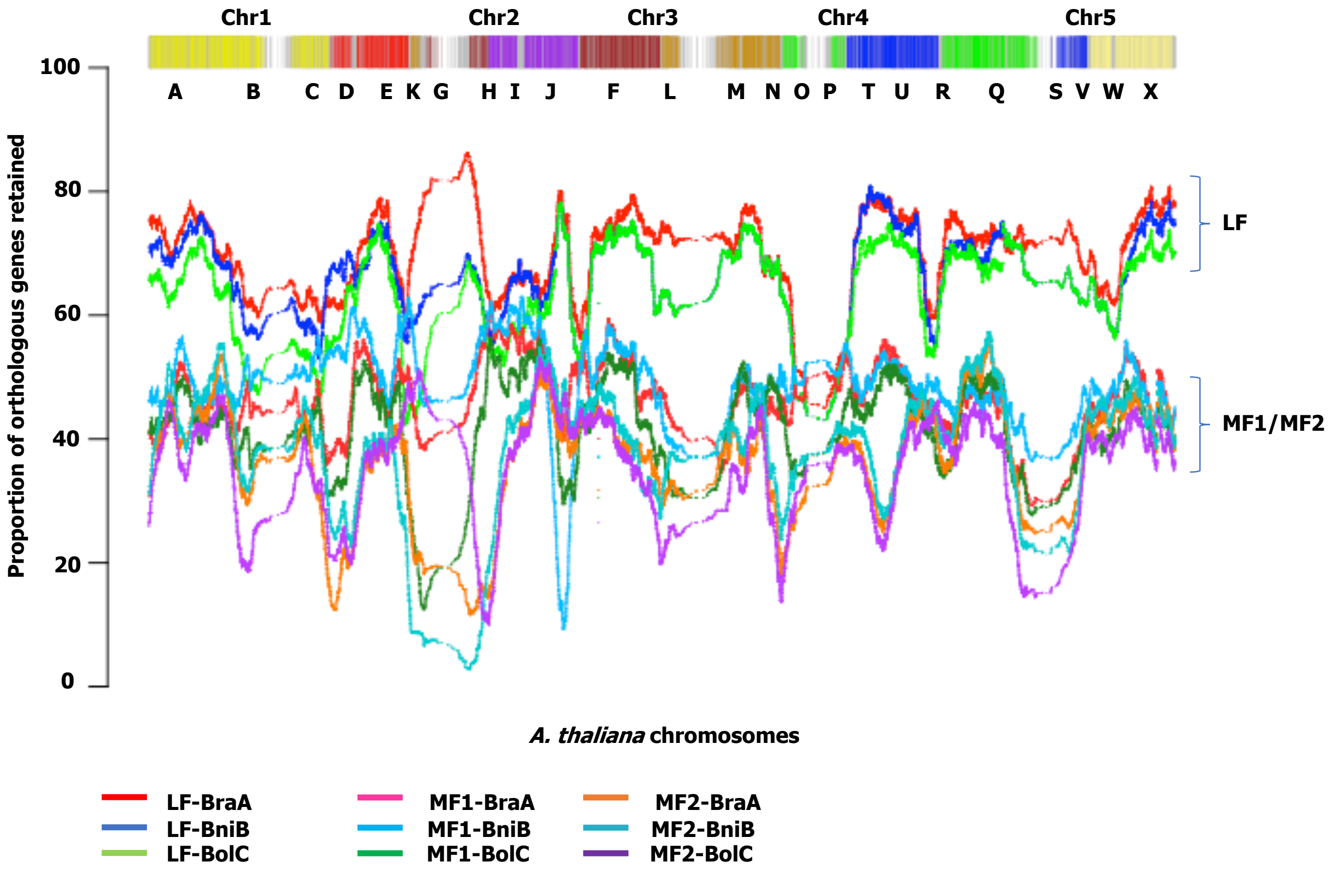


**Supplementary Fig. 6. Orthologous gene retention in the BraA[21], BniB (this study), and BolC[22] genomes corresponding to the *A. thaliana* genes.** Position of the *A. thaliana* genes have been plotted on axis X, the proportion of the genes retained in each of the three constituent paleogenomes of the A, B and C genomes has been plotted on axis Y. The constituent paleogenomes have been designated LF (least fragmented), MF1 (moderately fragmented), and MF2 (most fragmented) based on the percentage of genes retained in comparison to At, following the convention set for *B. rapa*[13].

V1- B01 – Lagercrantz[41]
V2- B02 - Punjabi et al.[11]
V3- B01 - New nomenclature

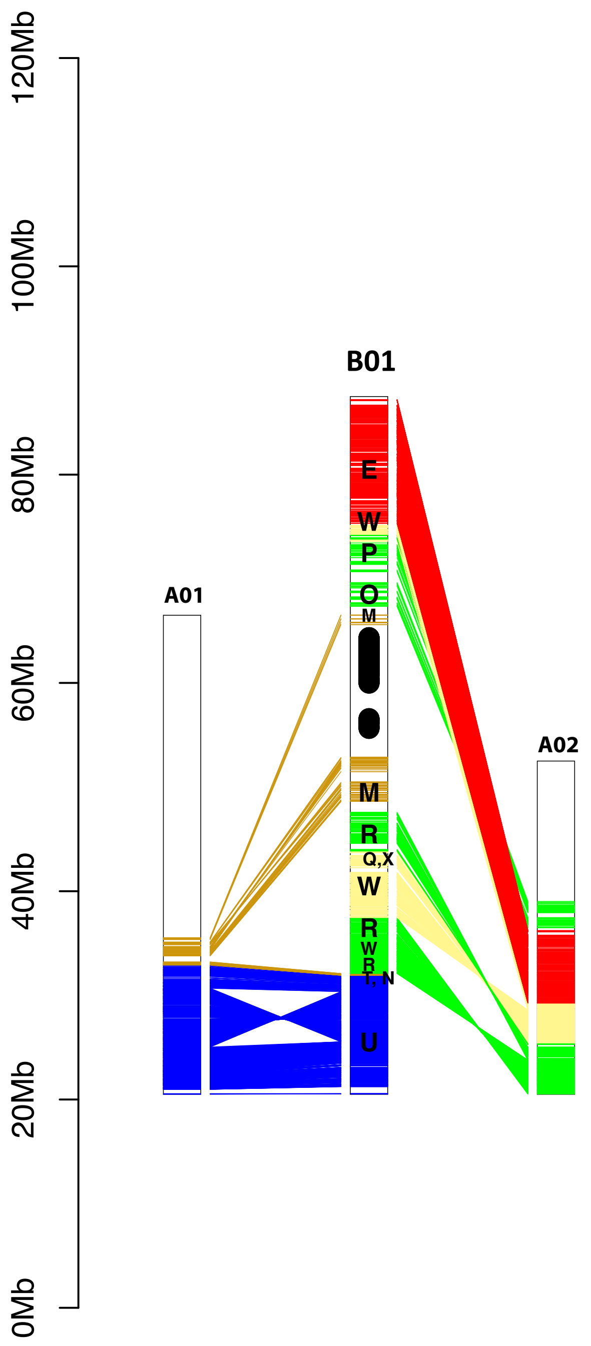


**Supplementary Fig. 7. Comparison of the eight *B. nigra* (BniB) pseudochromosomes with the ten *B. rapa* (BraA) pseudochromosomes[21] for homologous regions.** Each of the horizontal lines represents a gene. Homologous regions were identified by gene collinearity and the least Ks values amongst all the possible gene pairs. The number given to each B genome pseudochromosome in most of the cases is based on the number given to the A genome pseudochromosome with which it shows maximum homology.

V1- B06 - Lagercrantz[41]
V2- B06 - Punjabi et al.[11]
V3- B02 - New nomenclature

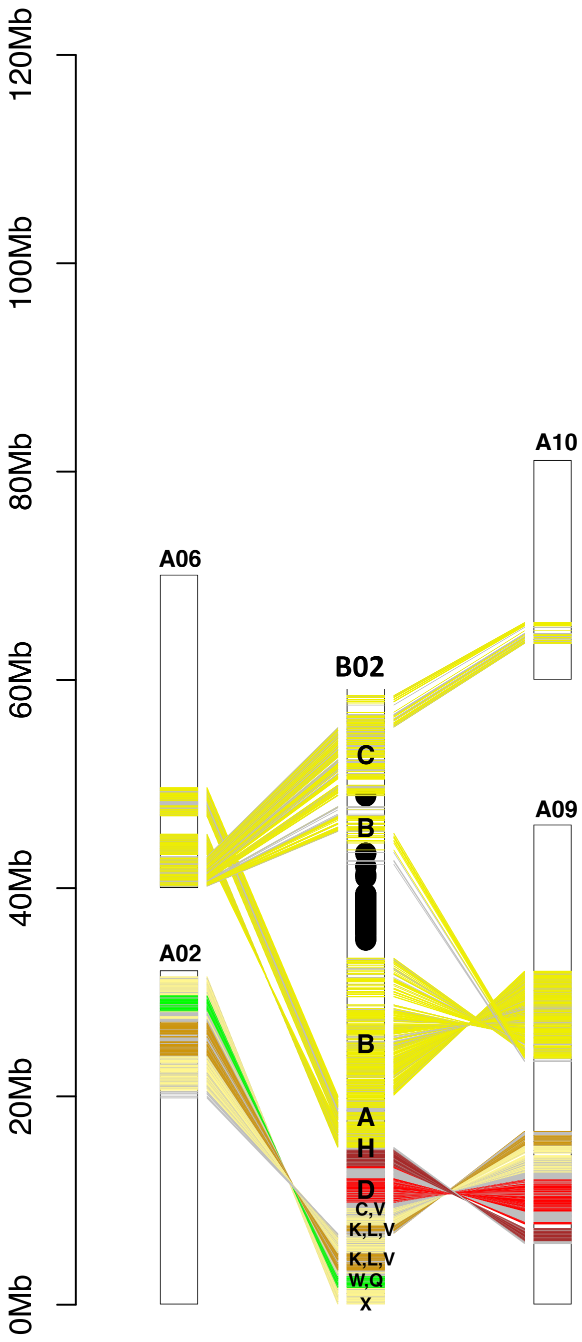

V1- B02 - Lagercrantz[41]
V2- B03 - Punjabi et al.[11]
V3- B03 - New nomenclature

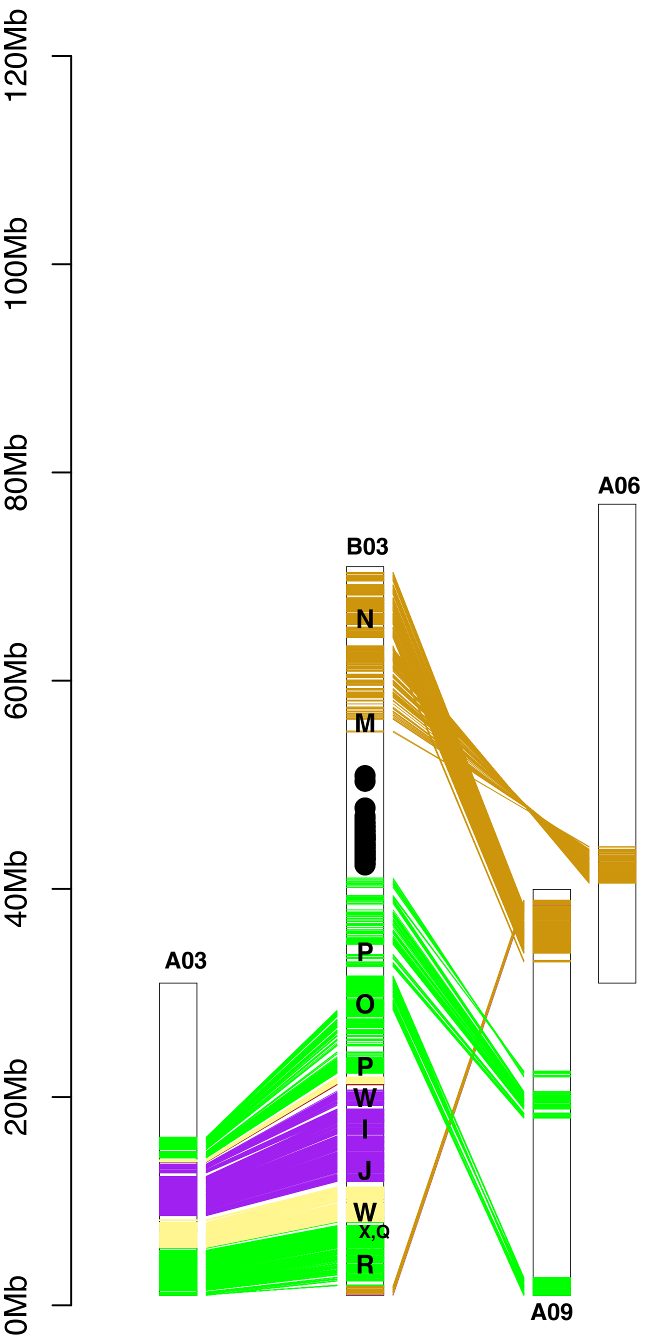

V1- B07 - Lagercrantz[41]
V2- B05 - Punjabi et al.[11]
V3- B04 - New nomenclature

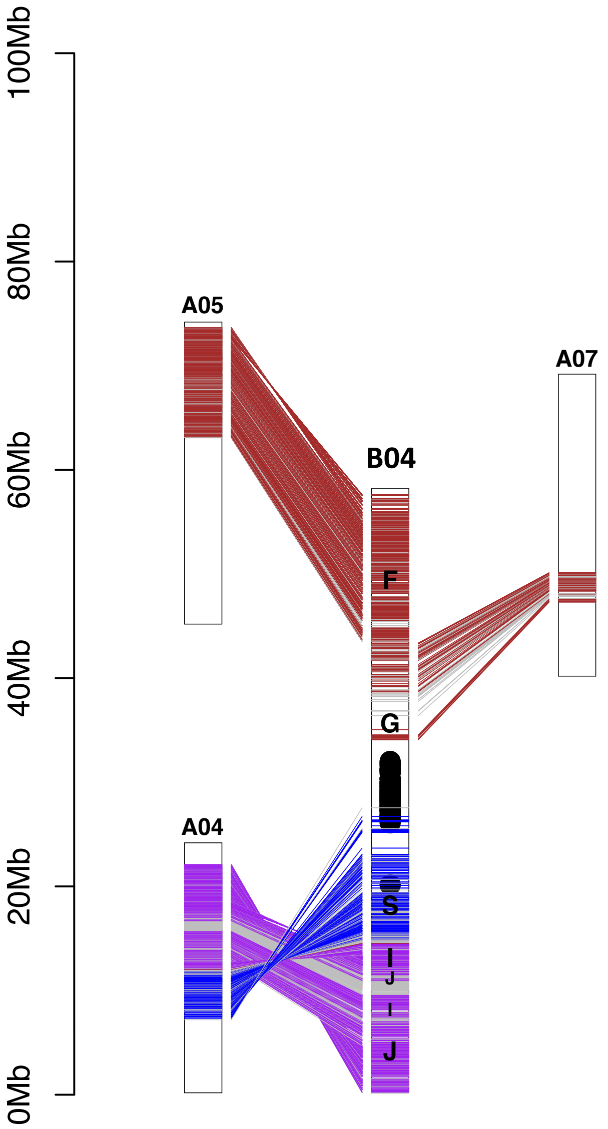

V1- B04 - Lagercrantz[41]
V2- B04 - Punjabi et al.[11]
V3- B05 - New nomenclature

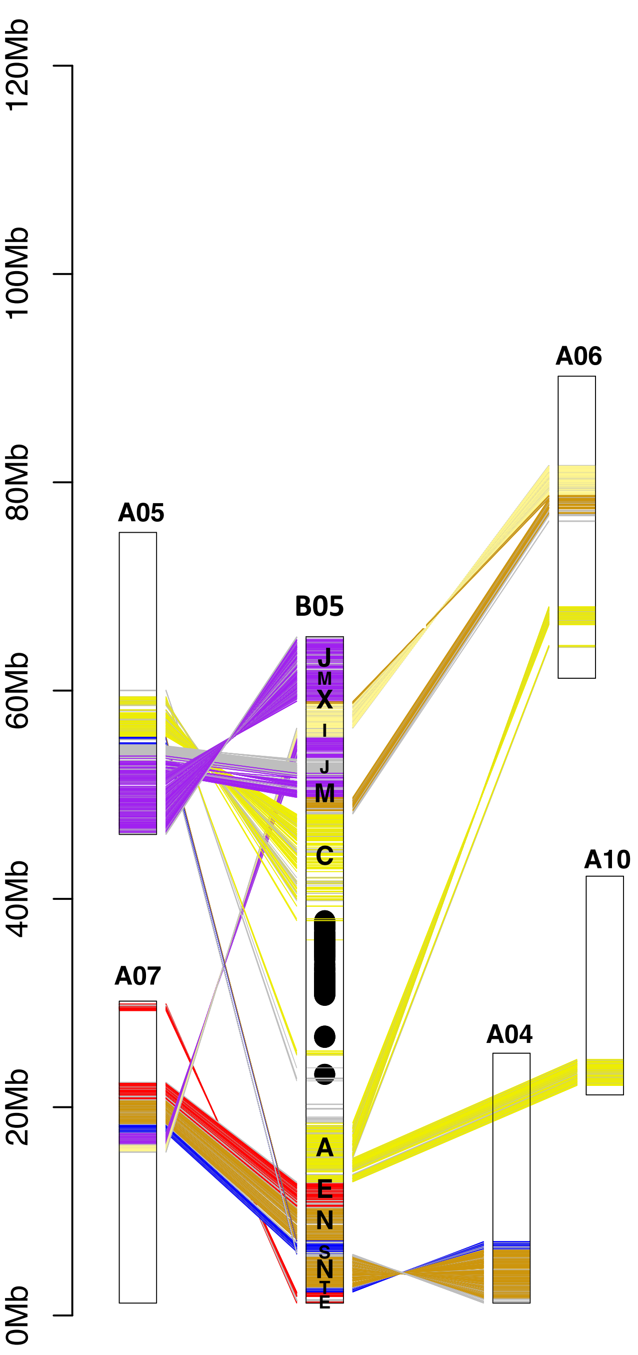

V1- B05 - Lagercrantz[41]
V2- B08 - Punjabi et al.[11]
V3- B06 - New nomenclature

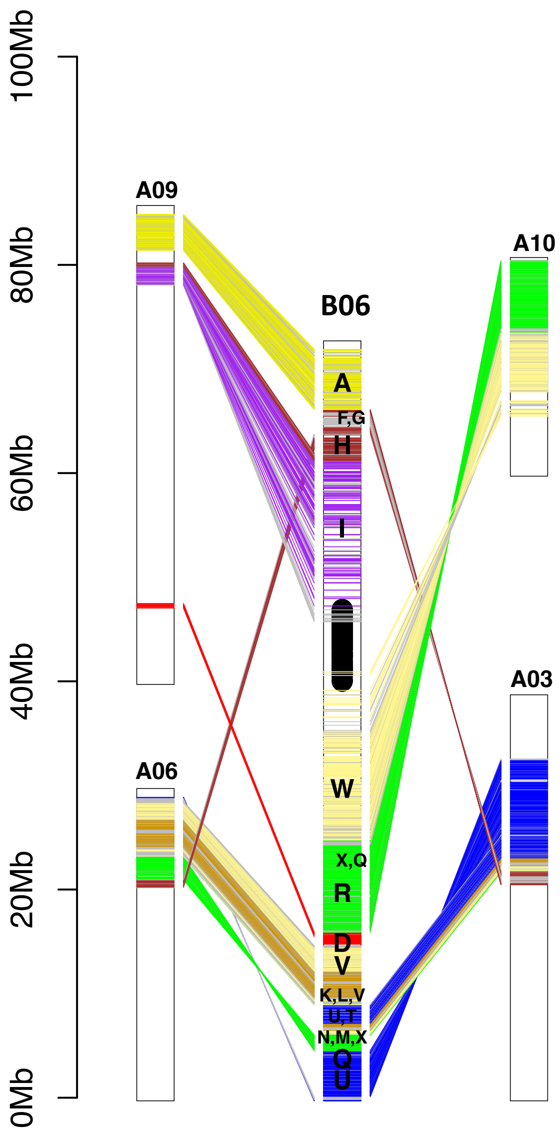

V1- B08 - Lagercrantz[41]
V2- B07 - Punjabi et al.[11]
V3- B07 - New nomenclature

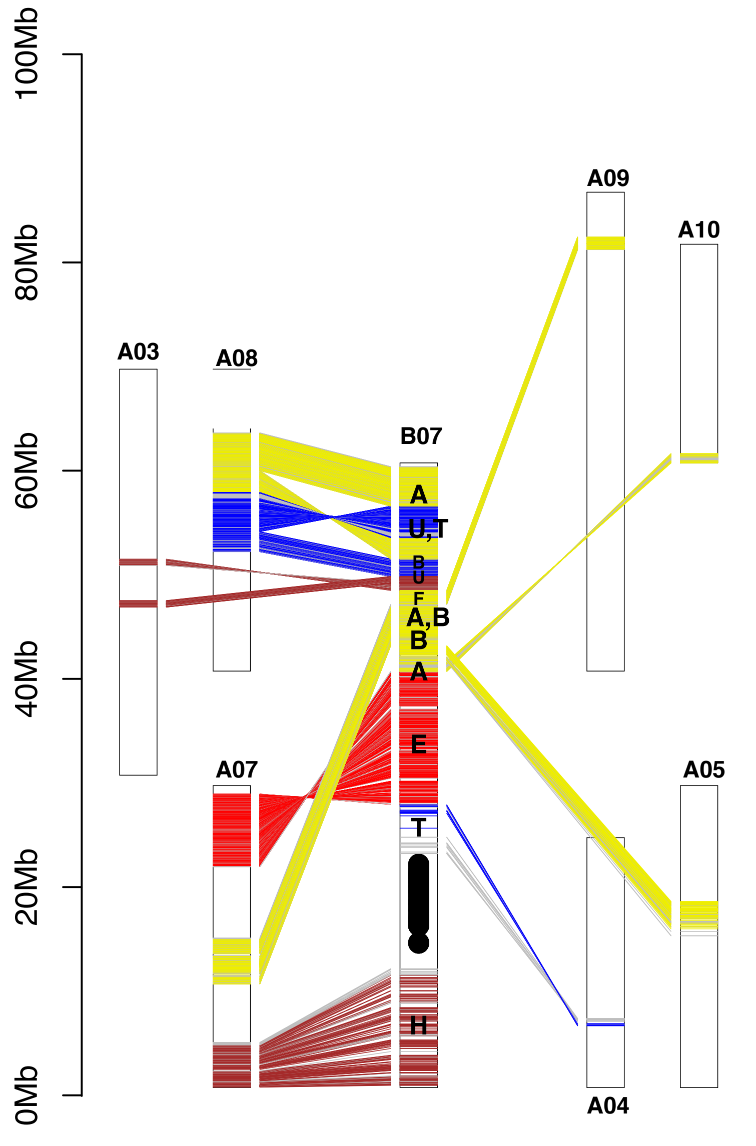

V1- B03 - Lagercrantz[41]
V2- B01 - Punjabi et al.[11]
V3- B08 - New nomenclature

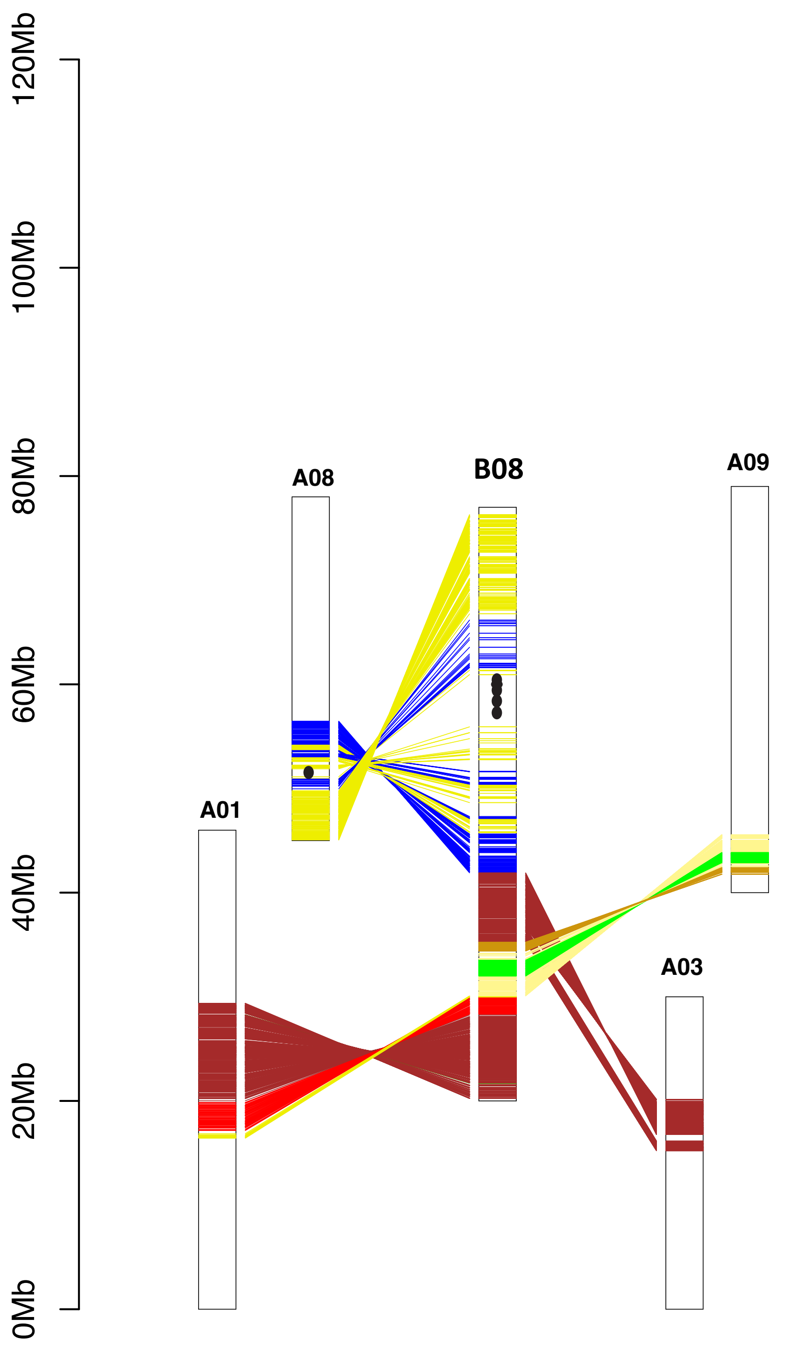
