## Supplementary File 1 B. nigra for "A highly contiguous genome assembly of *Brassica nigra* (BB) and revised nomenclature for the pseudochromosomes"

**Supplementary File 1 – RNA sequencing studies in *Brassica nigra***

Three different RNA sequencing analysis were carried out to validate the predicted genes in the *B. nigra* line Sangam (BnSDH-1) genome.

i. Illumina based transcriptome data taken from Paritosh et al.^23^ and mapped on the assembled BnSDH-1 genome.
ii. Illumina based transcriptome sequencing of *B. nigra* line 2782, carried out in the present study and mapped on the assembled BnSDH-1 genome.
iii. PacBio based transcriptome sequencing of *B. nigra* line BnSDH-1.

**Illumina assembly of *B. nigra* variety Sangam line BnSDH-1**

Illumina based transcriptome sequences of the line BnSDH-1 were taken from our previous study^23^. In brief, RNA from leaf, stem, and developing inflorescence tissues were pooled, and PE library was developed from 5µg of pooled RNA samples using the ‘mRNA library preparation kit’ (Illumina). The PE libraries with a mean insert size of ~250 bp were sequenced on Illumina Genome Analyzer IIx sequencer. A total of 117,766,948 raw sequence reads were produced. A detailed account of the parameters used for the raw-read filtering has been described in the Supplementary File of the Paritosh et al^23^. A total of 73,864,802 high-quality paired-end sequences of 70 bp length were obtained after the quality filtering step. These filtered PE reads were mapped on the assembled *B. nigra* BnSDH-1 genome with STAR aligner. The reads mapped on each gene were calculated and genes with more than 5 mapped reads were designated as expressed. A total of 30,901 genes were found to express in the transcriptome.

**Illumina based transcriptome assembly of line 2782**

For Illumina based transcriptome analysis of *B. nigra* line 2782, high-quality RNA was isolated from leaf, stem, and developing inflorescence tissues. Approximately 5µg of pooled RNA, with a RIN value 7.8, was used for library preparation following the method described earlier ^23^. PE library with mean insert size 278 bp was developed and sequenced on an Illumina Genome Analyzer IIx machine. The sequenced paired-end reads (2x 101 bp) were quality filtered with Trimmometic software and Fastx-Toolkit (<http://hannonlab.cshl.edu/fastx_toolkit/>). Parameters for the Trimmometic based QC was set as described earlier^23^. Reads with a filtered length of < 60 bp were removed from the analysis. The statistics of the transcriptome sequencing of *B. nigra* line 2782 are given in Supplementary Table S1.

**Supplementary Table S1 | Statistics of transcriptome sequencing of *B. nigra* line 2782**

| **Illumina raw reads** | **stats** |
| --- | --- |
| Library Type / Insert size | 278 bp |
| Total number of reads | 127,542,692 |
| Base trimmed paired-end reads |  |
| Clean pre-processed reads | 38,973,278 |
| % GC | 41.82 |
| % Q>30 | 97.45 |
| Data in Mb | 1038 |

The filtered sequences were mapped on the error corrected genome assembly of the line BnSDH-1 using STAR aligner. Reads mapped on each of the predicted genes were counted and genes with ≥5 mapped reads were considered as expressed. A total of 30,705 genes were identified to be expressed in the transcriptome sequencing.

**PacBio based transcriptome sequencing of the line BnSDH-1**

PacBio based transcriptome analysis was carried out to validate the predicted genes in the BnSDH-1 genome. The Iso-seq analysis also provided information on the full-length gene structures and isoforms of each of the predicted genes.

RNA was isolated from the pooled sample of the stem, leaf, and developing inflorescence tissues.

Three different libraries were developed, one each of size range 0.5-1 kb, 1-2 kb, and 2-6 kb. The libraries were sequenced separately using P6-C4 chemistry on the PacBio RSII system. In total, 54,303, 2,54,370 and 2,99,761 raw PacBio reads were generated for 0.5-1 kb, 1-2 kb and 2-6 kb libraries, respectively. Mean read of insert for size range 0.5-1 kb was 694 bp; for 1-2 kb and 2-6 kb groups the insert sizes were 1,151 bp and 1,467 bp, respectively. From the reads of different insert sizes, 25,534, 1,47,021 and 1,42,914 full length non-chimeric reads and 14,515, 86,694 and 1,45,603 partial reads were identified. The full-length non-chimeric sequences were used for clustering with the Iterative Clustering and Error Correction (ICE) algorithm and the non-full-length reads were used for the polishing of the ICE generated consensus sequence. Finally, clustering and polishing yielded 6,626, 28,698, and 17,198 full-length high-quality polished consensus sequences from the three libraries 0.5-1 kb, 1-2 kb, and 2-6 kb, respectively. ORF of the PacBio generated transcripts were predicted using ANGEL software (PacBio). A total of 24,986 low-quality isoforms were also assembled. The statistics of the PacBio based assembly are given in Table S2. The assembled sequences were mapped on the BnSDH-1 genome using the Minimap2 program. Out of 74,221 assembled isoforms, 13,618 isoforms were found to cover complete gene sequences; 691 genes were found to have more than one splice form.

**Table S2– Statistics of PacBio based transcriptome sequencing of *B. nigra***

|  | **Sangam 0.5-1 kb** | **Sangam 1-2 kb** | | **Sangam 2-6 kb** |
| --- | --- | --- | --- | --- |
| **Stats of Reads of Insert** |  | |  |  |
| Number of reads of insert | 54,303 | | 254,370 | 299,761 |
| Read Bases of Insert | 37,718,461 | | 292,854,895 | 439,986,819 |
| Mean Read Quality of Insert | 0.8948 | | 0.9332 | 0.9054 |
| Mean Read Length of Insert | 694 | | 1,151 | 1,467 |
| Mean Number of Passes | 10 | | 11 | 7 |
| **Classification of reads of insert** |  | |  |  |
| Number of three prime reads | 32,413 | | 186,465 | 192,234 |
| Number of five prime reads | 30,157 | | 171,802 | 187,094 |
| Number of poly-A reads | 32,679 | | 184,032 | 190,394 |
| Number of filtered short reads | 14,068 | | 19,663 | 10,806 |
| Number of full-length reads | 25,720 | | 148,013 | 143,352 |
| Number of non-full-length reads | 14,515 | | 86,694 | 145,603 |
| Number of full-length non-chimeric reads | 25,534 | | 147,021 | 142,914 |
| Average full-length non-chimeric read length | 779 | | 1036 | 1431 |
| **Cluster** |  | |  |  |
| Number of consensus isoforms | 11,181 | | 39,984 | 34,085 |
| Number of polished high-quality isoforms | 6,626 | | 28,698 | 17,198 |
| Number of polished low-quality isoforms | 4,555 | | 11,285 | 16,887 |
| Average consensus isoforms read length | 799 | | 1,108 | 1,527 |
